## Supplemental Figures S1-S7 for "Maternal DNMT3A-dependent de novo methylation of the zygotic paternal genome inhibits gene expression in the early embryo"

**Supplemental Tables and Figures**

**Supplemental Table 1** Processed data and lists of genes used to generate figures in this study.

**Supplemental Table 2** List of datasets mined and generated in this study

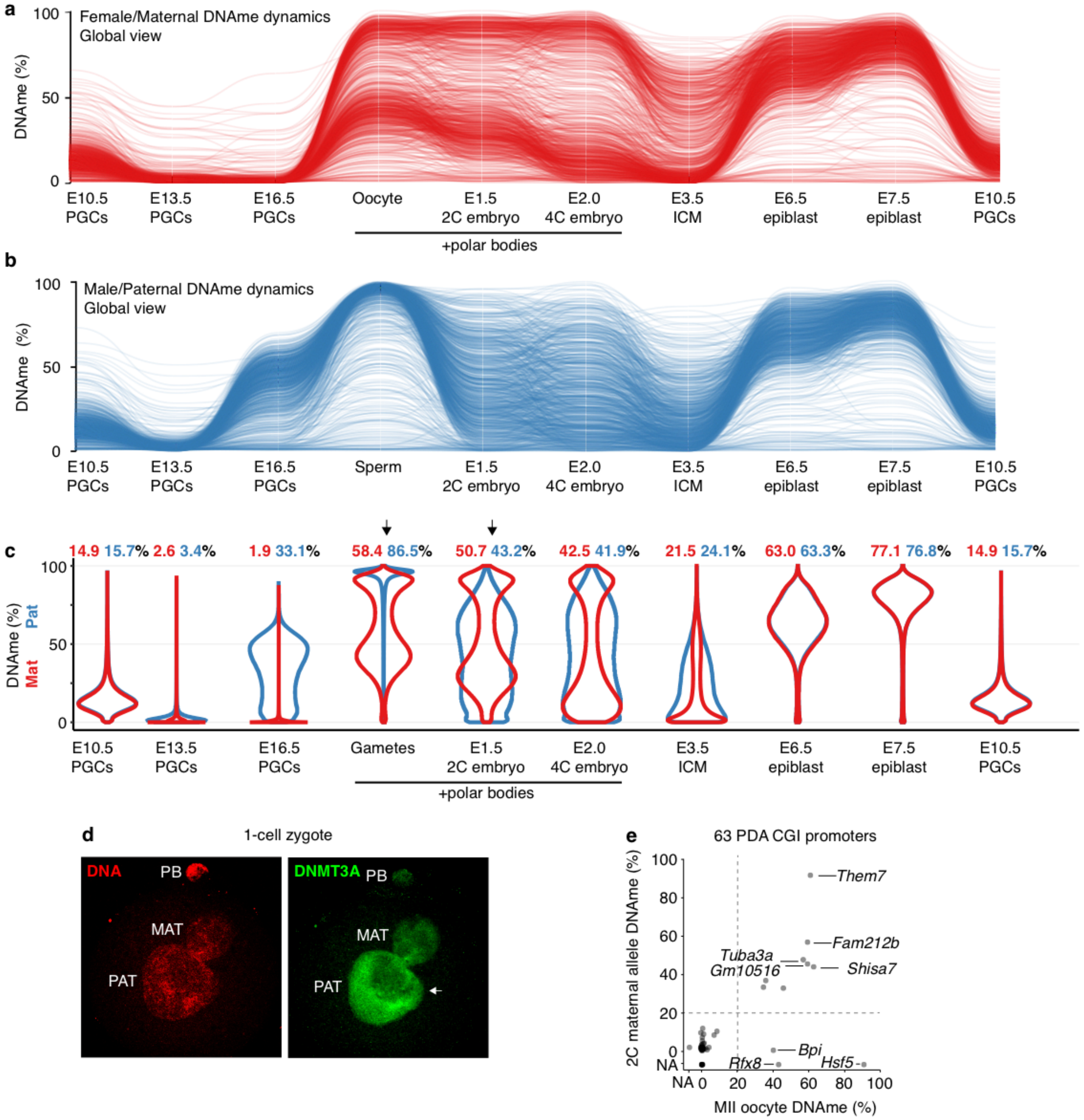


**Figure S1.** Female/maternal and male/paternal DNAme level dynamics during gametogenesis and embryonic development. **(a-b)** Parallel coordinate plots illustrating average DNAme levels over 2 kb genomic bins during **(a)** female/maternal and **(b)** male/paternal gametogenesis and embryonic development. Allele-specific analysis of WGBS data yielded 176,240 bins (roughly 15% of the mouse genome) overlapping at least 5 informative CpGs for which we could infer both maternal and paternal DNAme levels in all datasets. A random set of 1,000 bins on chromosome 19 are shown. MII oocyte, 2C and 4C embryo datasets include polar bodies. **(c)**

Distribution of global DNAme levels during gametogenesis and embryonic development. Mean DNAme percentages for each stage (red: female/maternal, blue: male/paternal) are shown above each violin plot. Arrows indicate the stages used to measure DNAme level change pre- and post-fertilization. **(d)** IF analysis of DNMT3A in a representative 1 cell zygote. Zygotic DNA is counterstained with propidium iodide (PI). PB; polar body. The white arrow indicates DNMT3A straining in the paternal pronucleus **(e)** Maternal allele DNAme levels over CGI promoters that show PDA (both datasets include PBs). NA: no data available for that particular stage.

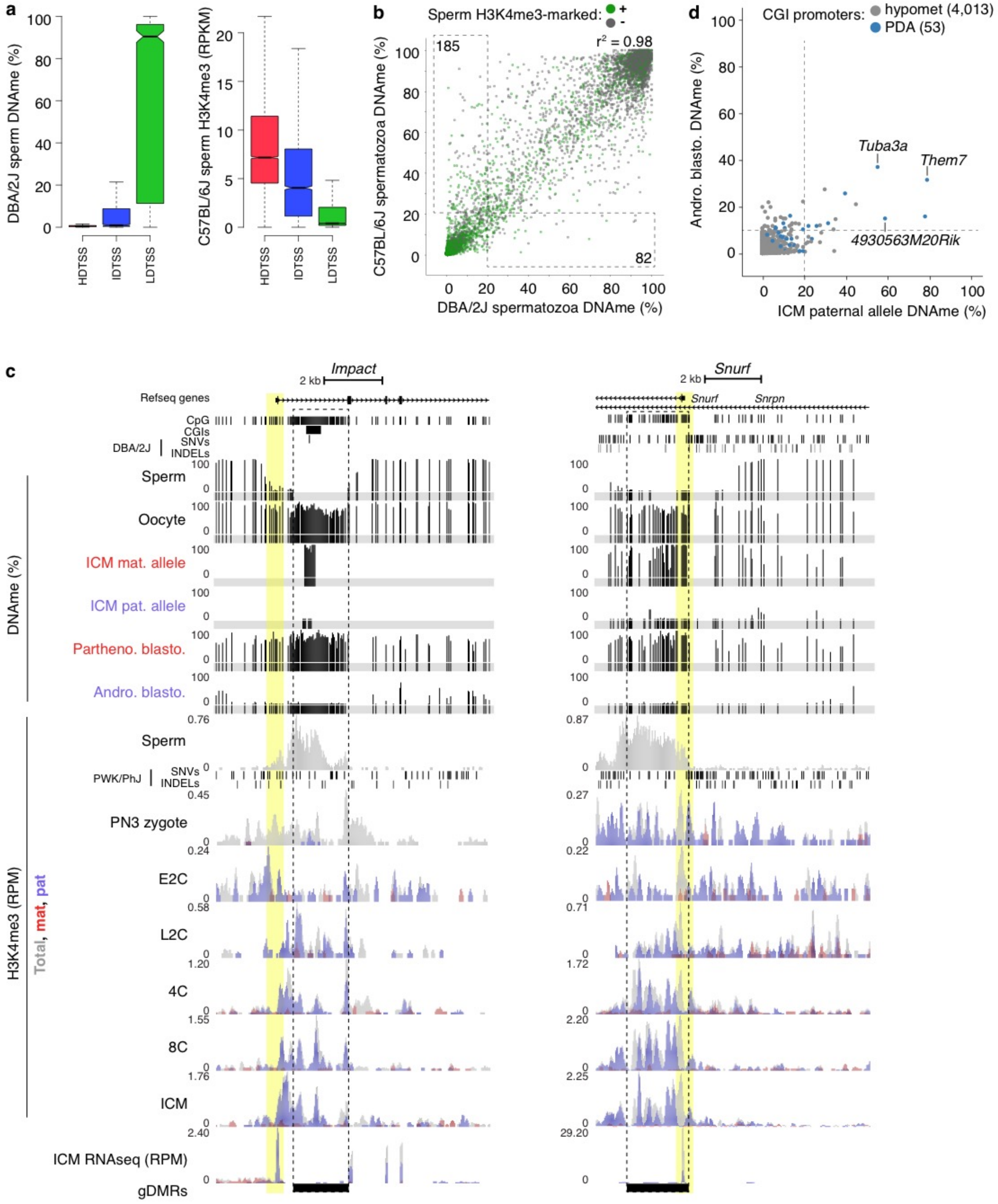


**Figure S2.** Defining hypomethylated CGI promoters in sperm. **(a)** Autosomal promoters (TSSs +/- 300 bp) were classified by CpG density as described previously[1], and the distribution of DNAme and H3K4me3 levels for each promoter category is shown. HD: high CpG density, ID: intermediate CpG density, LD: low CpG density. **(b)** 2D scatterplot illustrating the correlation of

DNAme levels between C57BL/6J and DBA/2J sperm. Data points are coloured by H3K4me3 enrichment in C57BL/6J sperm. Only promoters for which we calculated DNAme levels in both datasets (at least 2 CpGs with 5X coverage separated by >1 sequencing read length) are shown (n=20,583). 20,163 promoters had consistent DNAme levels (difference <20%) between strains. **(c)** UCSC genome browser screenshots of the *Impact* and *Snurf* paternally expressed imprinted genes. Promoters are highlighted in yellow, gametic DMRs by a dashed box and the location of informative CpGs (5X coverage) for each WGBS dataset are highlighted in grey. Embryonic H3K4me3 and ICM RNA-seq data are represented as a composite track containing total (allele-agnostic, grey), maternal (red) and paternal (blue) genomic tracks. NCBI Refseq genes, all CpG dinucleotides, CpG islands and genetic variants used in our allele-specific analyses (SNVs and INDELs) are also included. Mat.: maternal, Pat.: paternal, Partheno.: parthenogenetic, Andro.: androgenetic, Blasto.: blastocyst, PN3: pronuclear stage 3, E2C: early 2 cell, L2C: late 2 cell, 4C: 4 cell, 8C: 8 cell, ICM: inner cell mass cells. **(d)** Scatterplot showing the association between CGI promoter DNAme levels on the paternal allele of normal F1 hybrid ICM cells and androgenetic blastocysts.

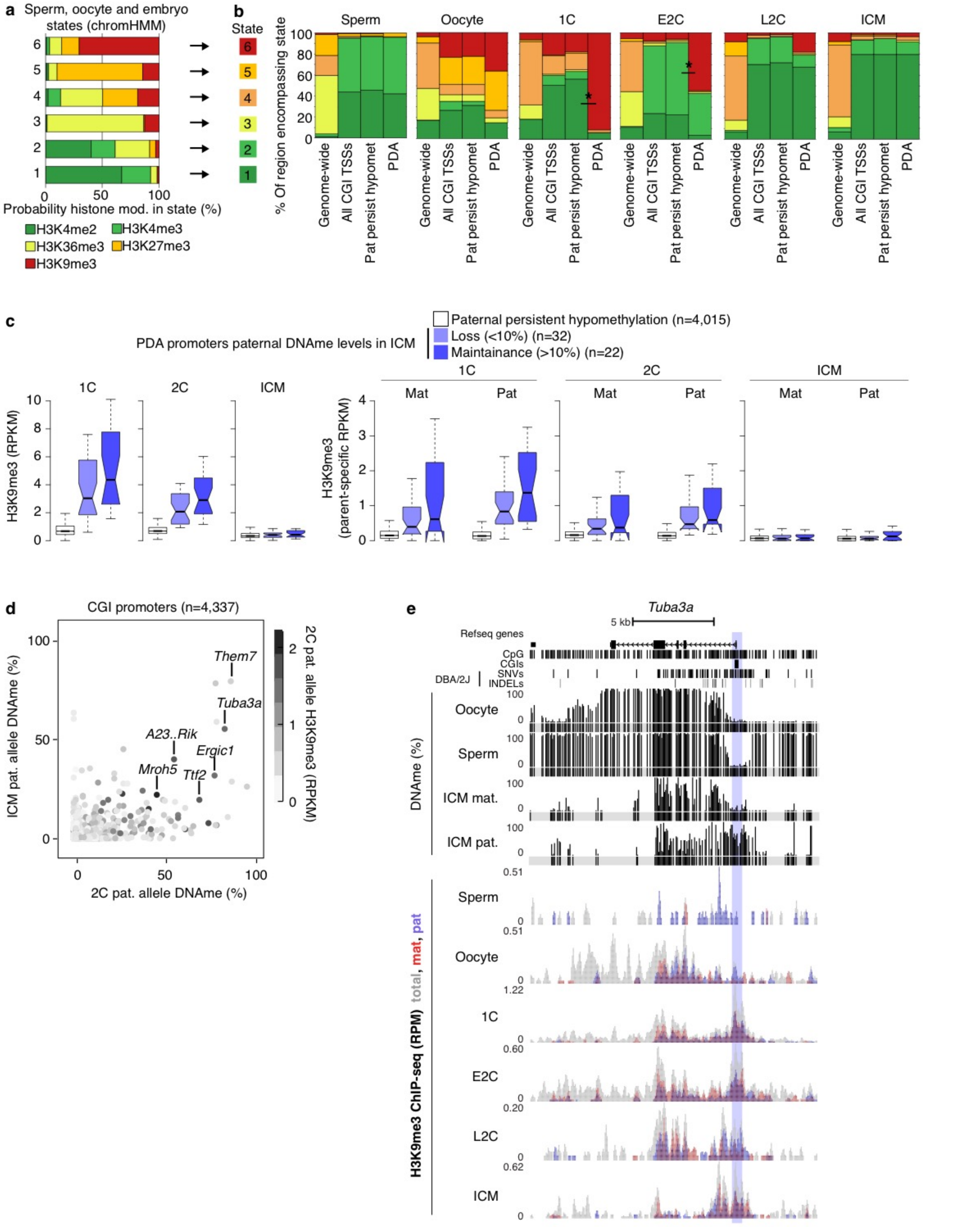


**Figure S3.** Relationship between histone PTMs and PDA **(a)** Six distinct chromatin states were identified using ChromHMM and ChIP-seq data (H3K4me2, H3K4me3, H3K36me3, H3K27me3 and H3K9me3) from spermatozoa, MII oocytes, 1C, early 2C, late 2C and blastocyst-stage embryos. The probability that a histone modification is found within each state is shown. Based on these probabilities, a colour code was assigned to each state (right). **(b)** The relative enrichment of each chromatin state genome-wide and over genomic regions of interest, including all autosomal CGI promoters (n=13,342), those that show persistent paternal DNA hypomethylation following fertilization (n=4,315) and those that show PDA (n=63), is shown. **(c)** The distribution of total (left), and parent allele-specific H3K9me3 (right) levels over CGI promoters in 1C, E2C and blastocyst-stage embryos is shown. CGI promoters were categorized by paternal DNAme dynamics, including persistent hypomethylation following fertilization (n=4,015), PDA genes showing loss of paternal DNAme by the blastocyst stage (n=32), and PDA genes showing persistence of paternal DNAme in the blastocyst (n=22). **(d)** 2D scatterplot illustrating the association between H3K9me3 and DNAme levels on the paternal allele at the 2C stage and maintenance of DNAme on the paternal allele in ICM cells. **(e)** UCSC genome browser screenshot of the *Tuba3a* locus illustrating DNAme levels in gametes and the ICM as well as maternal (red) and paternal (blue) H3K9me3 levels. The CpG rich promoter of *Tuba3a* is highlighted in blue and the location of informative CpGs (5X coverage) for each WGBS dataset are highlighted in grey. The genomic locations for NCBI Refseq genes, all CpG dinucleotides, CpG islands and genetic variants used in our allele-specific analyses (SNVs and INDELs) are also shown. H3K9me3 ChIP-seq data are represented as a composite tracks containing total (allele-agnostic, grey), maternal (red) and paternal (blue) genomic tracks.

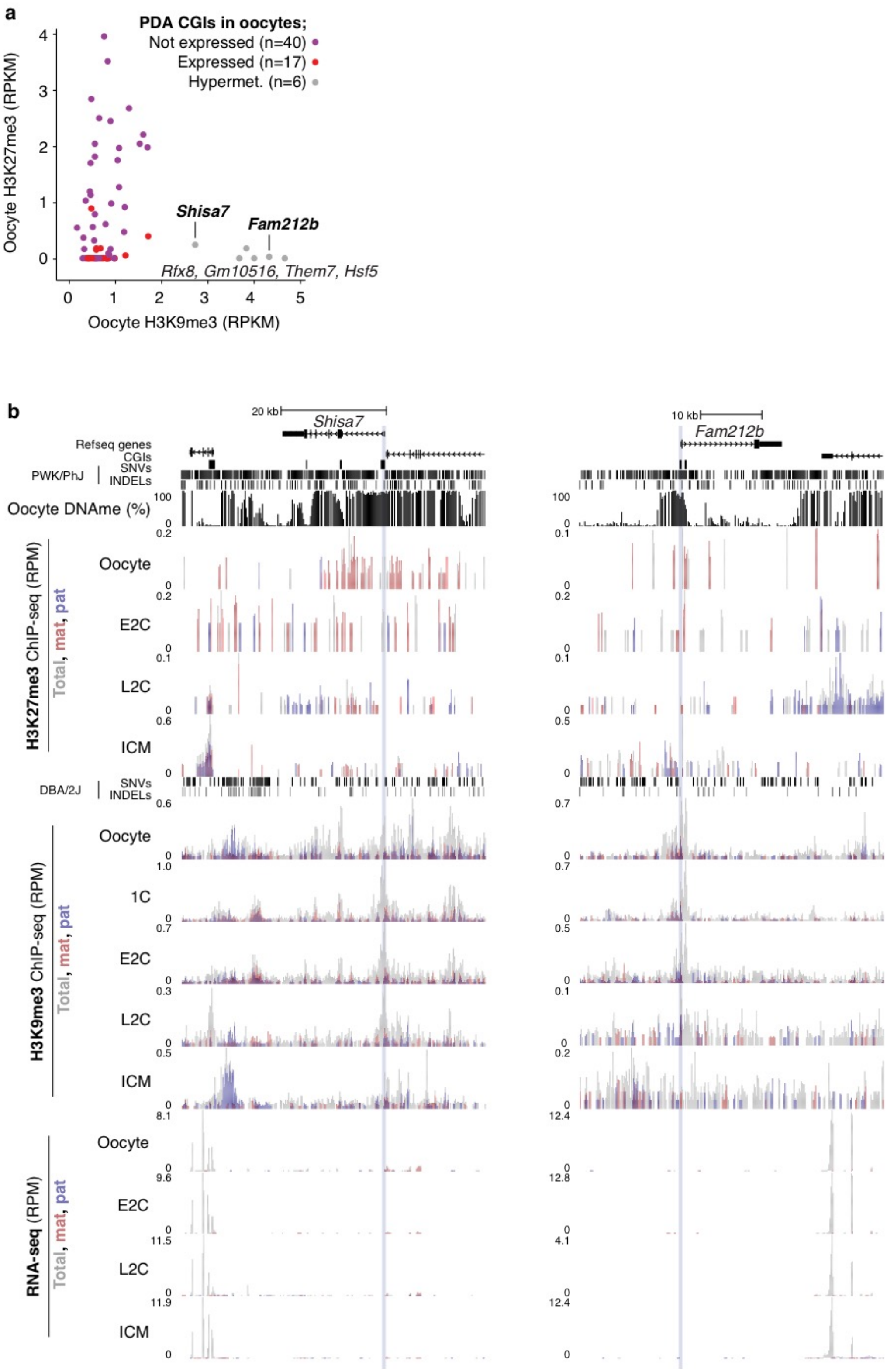


**Figure S4** Transcriptionally silenced PDA CGI promoters in oocytes are either DNA hypermethylated and H3K9me3-marked or hypomethylated and H3K27me3-marked in oocytes and the early embryo. **(a)** Scatterplot showing H3K9me3 and K3K27me3 enrichment in MII oocytes. Data points are coloured as in Fig. 3b. **(b)** UCSC genome browser screenshots of the transcriptionally silenced *Shisa7* and *Fam212b* loci illustrating hypermethylation and H3K9me3-enrichment of their CGI promoters in oocytes. Allele-specific H3K27me3, H3K9me3 and expression data from oocytes and the early embryo are included. 1C: 1 cell zygote, E2C: early 2 cell, L2C: late 2 cell, ICM: inner cell mass cells.

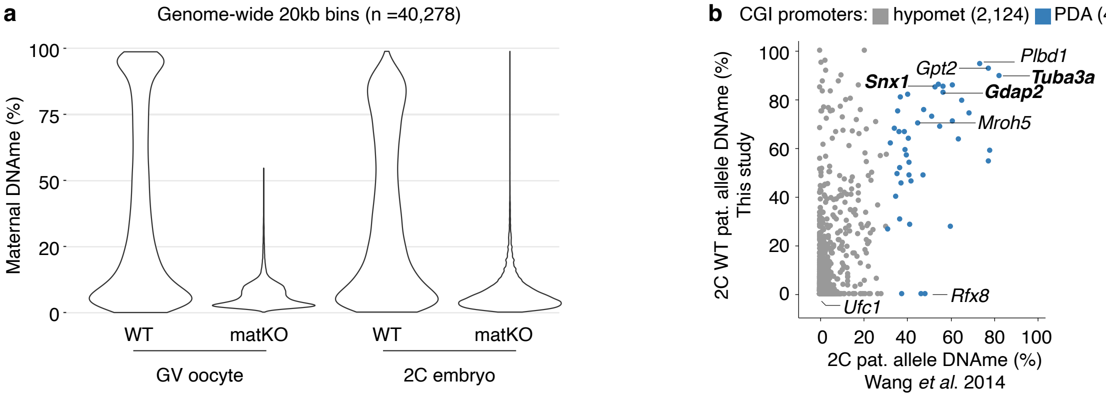


**Figure S5.** DNAme analysis of maternal *Dnmt3a* KO embryos **(a)** Distribution of maternal DNAme levels over 20kbp genomic bins in WT and *Dnmt3a* matKO GV oocytes and 2 cell embryos. 20kb bins overlapping >3 informative CpGs (covered by 5X reads and separated by >1 sequencing read length) in all 4 datasets are reported (n=40,278). GV oocyte data from Shirane et al. 2013. **(b)** Paternal allele CGI promoter DNAme level association between wild-type 2C data generated in this study and those from[2]. Genes are coloured by paternal DNAme dynamics immediately following fertilization: grey; persistent hypomethylation on the paternal genome (paternal hypomet, n=2,124), blue; paternal DNAme acquisition (PDA, n=43). Paternal DNAme levels over CGI promoters were reported if they were covered by at least 5 informative CpGs (covered by 1X read and separated by >1 sequencing read length) in our 2C WGBS datasets.

Richard_Albert_FigS6

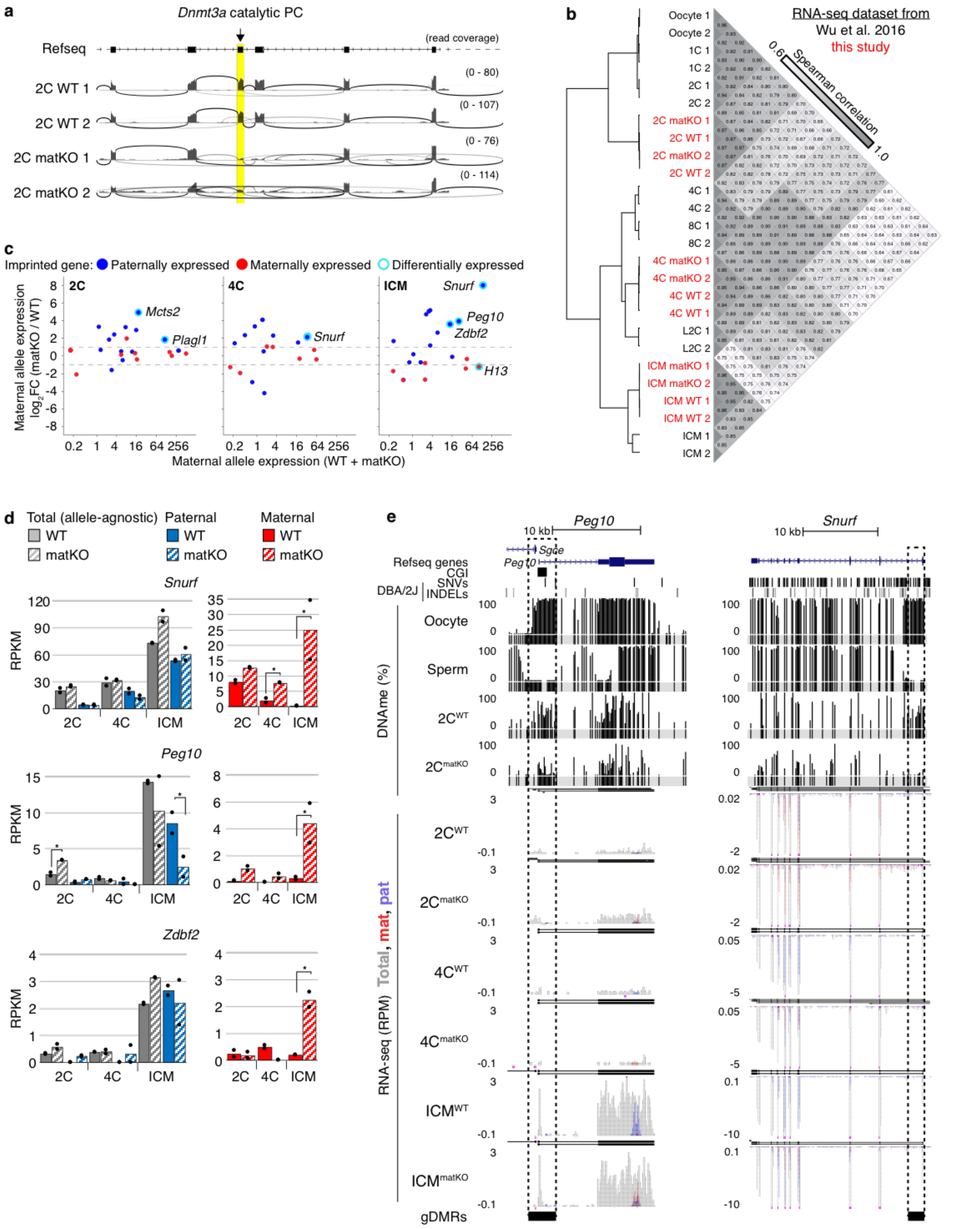


**Figure S6.** Transcriptomic analysis of maternal *Dnmt3a* KO embryos. **(a)** 2 cell strand-specific RNA-seq sashimi plot illustrating deletion of the catalytic exon of *Dnmt3a*. **(b)** Spearman correlation of autosomal gene expression (log2(RPKM+1) in datasets mined from[3] (black) and generated in this study (red). **(c)** 2D scatterplots showing change in known imprinted gene expression from the maternal allele. Data points are coloured by whether they are normally paternally (blue) or maternally (red) expressed in somatic cells. Differentially expressed (≥2-fold change, Benjamini-Hochberg adjusted P-value ≤0.1) genes are highlighted in teal. **(d)** Bar plots illustrating differential expression of select imprinted genes from both alleles (total, grey), the paternal genome (blue) and the maternal genome (red) in wild-type (filled bars) and *Dnmt3a* matKO (striped bars) embryos. Each bar represents the mean expression value of two biological replicates (dots) in RPKM. Statistically significant differential expression is indicated by an asterisk. **(e)** UCSC genome browser screenshots of the paternally expressed imprinted genes *Peg10* and *Snurf* illustrating loss of imprinting. Gametic DMRs are indicated by a dashed box and the location of informative CpGs (5X coverage) for each WGBS dataset are highlighted in grey. Strand-specific RNA-seq data is represented as a composite track of biological duplicates containing total (allele-agnostic, grey), maternal (red) and paternal (blue) genomic tracks. *De novo* assembly of transcripts is included above each RNAseq dataset. NCBI Refseq genes, all CpG dinucleotides, CpG islands and genetic variants used in our allele-specific analyses (SNVs and INDELs) are also included.

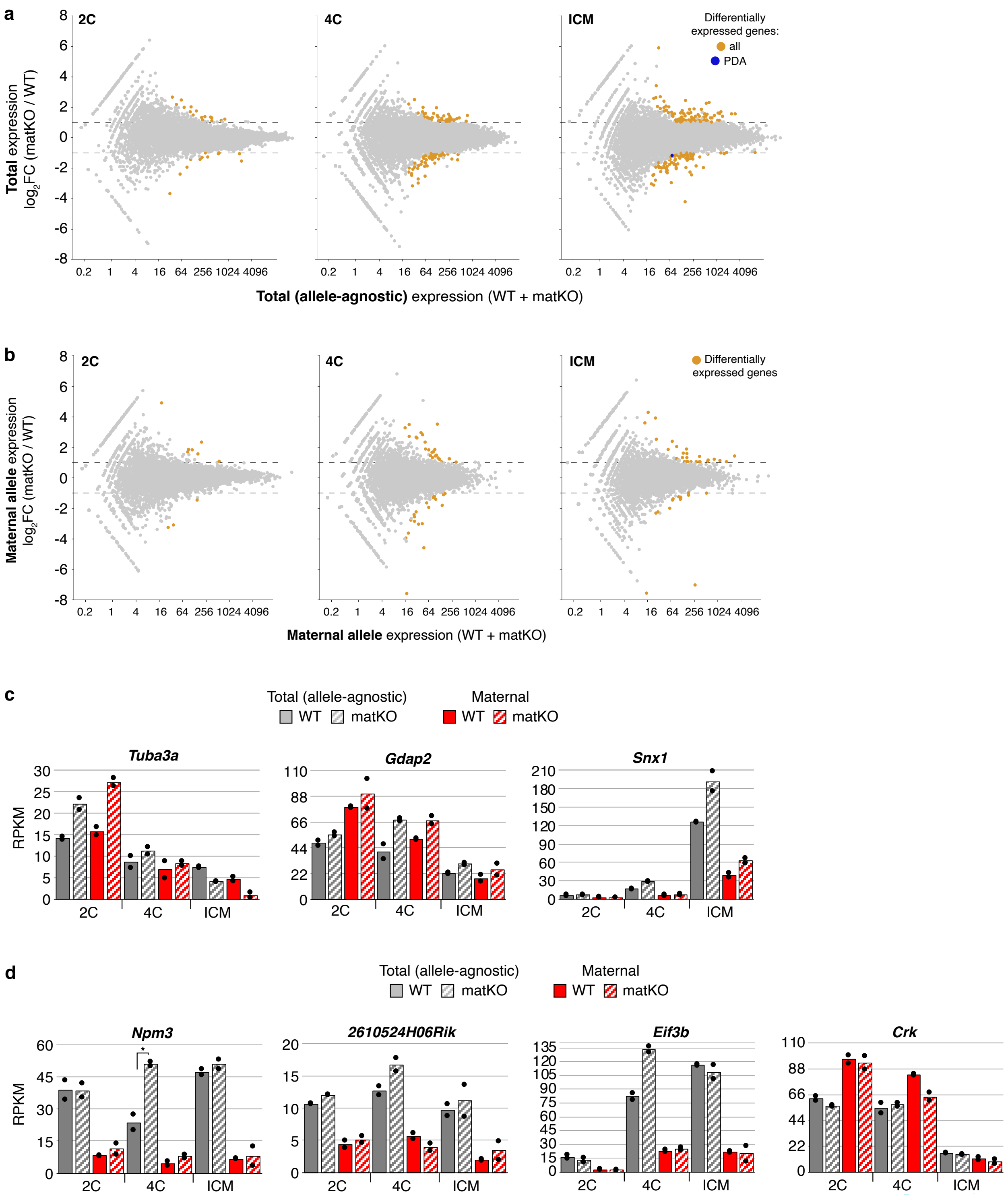


**Figure S7.** Allele-agnostic and maternal-allele analysis of the change in CGI promoter gene expression in *Dnmt3a* matKO embryos. **(a)** 2D scatterplots showing average (x-axis) and differential (y-axis) total (allele-agnostic) expression between wild-type WT and matKO 2C, 4C embryos and ICM cells. Genes that show statistically significant differences in expression (≥2-

fold change, Benjamini-Hochberg adjusted P-value ≤0.1) are highlighted in orange, and those that show PDA are coloured in blue. **(b)** 2D scatterplots showing average (x-axis) and differential (y-axis) maternal-allele expression between wild-type WT and matKO 2C, 4C embryos and ICM cells. Legend as in **a**. No genes that show PDA at their CGI promoters were statistically differentially expressed from the maternal allele. **(c-d)** Bar charts illustrating differential expression of select genes from both alleles (grey) and the maternal genome (red) in wild-type (filled bars) and *Dnmt3a* matKO (striped bars) embryos. Each bar represents the mean expression value of two biological replicates (dots) in RPKM. Statistically significant differences (Benjamini-Hochberg adjusted P value ≤0.1) are indicated by an asterisk.

**Supplemental Figure References**


1. Weber, M. *et al.* Distribution, silencing potential and evolutionary impact of promoter DNA methylation in the human genome. *Nat Genet* **39,** 457–466 (2007).
2. Wang, L. *et al.* Programming and inheritance of parental DNA methylomes in mammals. *Cell* **15,** 979–991 (2014).
3. Wu, J. *et al.* The landscape of accessible chromatin in mammalian preimplantation embryos. *Nature* **534,** 652–657 (2016).
